# The Sec1/Munc18 (SM) protein Vps45 is involved in iron uptake, mitochondrial function and virulence in the pathogenic fungus *Cryptococcus neoformans.*

**DOI:** 10.1101/356535

**Authors:** Mélissa Caza, Guanggan Hu, Eric David Neilson, Minsu Cho, Won Hee Jung, James W. Kronstad

## Abstract

The battle for iron between invading microorganisms and mammalian hosts is a pivotal determinant of the outcome of infection. The pathogenic fungus, *Cryptococcus neoformans*, employs multiple mechanisms to compete for iron during cryptococcosis, a disease primarily of immunocompromised hosts. In this study, we examined the role of endocytic trafficking in iron uptake by characterizing a mutant defective in the Sec1/Munc18 (SM) protein Vps45. This protein is known to regulate the machinery for vesicle trafficking and fusion via interactions with SNARE proteins. As expected, a *vps45* deletion mutant was impaired in endocytosis and showed sensitivity to trafficking inhibitors. The mutant also showed poor growth on iron-limited media and a defect in transporting the Cfo1 ferroxidase of the high-affinity iron uptake system from the plasma membrane to the vacuole. Remarkably, we made the novel observation that Vps45 also contributes to mitochondrial function in that a Vps45-Gfp fusion protein associated with mitotracker, and a *vps45* mutant showed enhanced sensitivity to inhibitors of electron transport complexes as well as changes in mitochondrial membrane potential. Consistent with mitochondrial function, the *vps45* mutant was impaired in calcium homeostasis. To assess the relevance of these defects for virulence, we examined cell surface properties of the *vps45* mutant and found increased sensitivity to agents that challenge cell wall integrity and antifungal drugs. A change in cell wall properties was consistent with our observation of altered capsule polysaccharide attachment, and with attenuated virulence in a mouse model of cryptococcosis. Overall, our studies reveal a novel role for Vps45-mediated trafficking for iron uptake, mitochondrial function and virulence.

## AUTHOR SUMMARY

*Cryptococcus neoformans* is a causative agent of cryptococcal meningitis, a disease that is estimated to cause ~ 15% of AIDS-related deaths. In this context, cryptococosis is the second most common cause of mortality in people with HIV/AIDS, closely behind tuberculosis. Unfortunately, very few antifungal drugs are available to treat this disease. However, understanding mechanisms involved in the pathogenesis of *C. neoformans* can lead to new therapeutic avenues. In this study, we discovered a new role for a regulatory protein involved in vesicle transport. Specifically, we found that the Vps45 protein, which regulates vesicle fusion, participates in the trafficking of iron into fungal cells, supports mitochondria function, mediates antifungal resistance and is required for virulence. These discoveries shed light on the molecular mechanisms underlying the uptake and use of iron as an essential nutrient for the virulence of *C. neoformans.* Further investigations could lead to the development of drugs that target Vps45-mediated processes.

## INTRODUCTION

The pathogenic fungus *Cryptococcus neoformans* attacks immunocompromised people to cause cryptococcosis, a particularly devastating disease in HIV/AIDS sufferers (1). Adaptations of the fungus to cause disease in mammalian hosts include the ability to grow at 37°C, to deliver key virulence components to the external milieu, and to acquire nutrients for proliferation (2,3). In the latter case, iron plays a key role in the virulence of *C. neoformans* as a cofactor in essential biochemical reactions and as a regulator of the elaboration of the polysaccharide capsule, a major virulence factor (4,5). As with other pathogens, *C. neoformans* must compete against host nutritional immunity to obtain iron during infection. Iron withholding by the host is achieved by iron-binding proteins such as transferrin, lactoferrin, and ferritin that maintain available iron at extremely low levels (6,7). On the other hand, iron overload exacerbates cryptococcal disease in a mouse model of cryptococcosis (8). Because of the iron-limited nature in the host, *C. neoformans* has developed multiple strategies to acquire iron including the use of a high-affinity iron uptake system composed of the cell surface iron permease Cft1 and the ferroxidase Cfo1 (9,10), the secreted mannoprotein Cig1 for iron uptake from heme (11) and the requirement of the endosomal sorting complex required for transport (ESCRT) pathway for endocytosis and intracellular trafficking of exogenous heme (12,13). Furthermore, these systems are known to participate in the virulence of *C. neoformans* in a murine inhalation model of cryptococcosis (9–13).

The discovery that the ESCRT pathway is critical for iron acquisition from exogenous heme implies that intracellular transport occurs via endocytosis. Endocytic pathways internalize extracellular components, fluids and membrane-bound factors, including lipids and proteins into cytoplasmic vesicles. These cargo-loaded vesicles fuse with the early endosome (EE) where sorting by ESCRT machinery and maturation occurs into late endosomes (LE) or multivesicular bodies (MVBs). Depending on its nature, cargo is either sent to the vacuole for degradation, to other organelles, or it may be recycled back to the trans-Golgi network (TGN) where re-secretion to the plasma membrane may occur (14–16). The hypothesis that endocytic processes are involved in iron uptake in fungi was first inferred from studies in *Saccharomyces cerevisiae* and *Candida albicans.* For example, the ferrichrome siderophore receptor Arn1p in *S. cerevisiae* is found in endosome-like vesicles and is sorted directly from the Golgi to the endosomal compartment in the absence of its substrate. However, when cells are exposed to ferrichrome, Arn1p is relocalized to the plasma membrane where the siderophore-loaded receptor complex rapidly undergoes endocytosis (17). In *C. albicans,* the plasma membranes proteins Rbt5 and Rbt51 are required for binding and iron uptake from heme and hemoglobin. A screen for mutants defective in iron utilisation from hemoglobin in *S. cerevisiae* expressing heterologously Rbt51 revealed mutants impaired in vacuolar functions, including vacuolar ATPase, and components of the ESCRT pathway and the HOPS complex. Subsequent analysis confirmed the role of ESCRT-I complex *VPS23* and *VPS28* in iron uptake from hemoglobin, but not from ferrichrome, suggesting that the processes of iron uptake from hemoglobin and ferrichrome are distinct (18). Likewise, ESCRT mutants in *C. neoformans* show defects in iron acquisition from heme and hemoglobin, but not from ferrichrome (12,13). In addition, there was an extended lag phase for ESCRT mutants when grown in media containing iron chloride as a sole iron source suggesting that ESCRT pathway plays a role in recycling the high-affinity iron uptake system Cft1-Cfo1 to the plasma membrane (9,10). This idea is consistent with the findings in *S. cerevisiae* that levels of iron influence cellular localization of the iron permease/ferroxidase proteins Ftr1 and Fet3 (i.e., recycling back to the plasma membrane in low levels of iron or targeting to the vacuole for degradation in high levels of iron); these endocytic sorting processes also require the ESCRT pathway (19).

Vesicle fusion events in endocytic compartments are a dynamic process that is regulated by specific mediators such as tethering factors, SNAREs, Rab GTPases, guanine-nucleotide exchange factors (GEFs) and Sec1/Munc18 (SM) proteins. SNARE (soluble N-ethylmaleimide-sensitive attachment protein receptor) proteins form tight four-helix bundles (core complexes) that bring specific membranes together (20). Prior to fusion, transport vesicles are loaded with tethering factors that promote the assembly of specific and functional SNARE complexes (also called SNAREpins) by interacting with SM proteins, Rab GTPases and GEFs (21). SM proteins are evolutionarily conserved proteins that are considered to be key elements of the fusion machinery because they strongly promote SNARE-mediated fusion and specificity (22,23). Four SM proteins were identified in *S. cerevisiae* and they function at distinct intracellular transport steps depending on their cognate partners. For example, the SM protein Sec1 functions at the plasma membrane during exocytosis while Sly1 operates at the endoplasmic reticulum (ER)-Golgi membrane (21). The SM protein Vps33 is required for delivery of endocytosed cargo into the vacuole. Vps33 is a core member of the endosomal Class C core vacuole/endosome transport (CORVET) complex and the vacuolar homotypic fusion and vacuole protein sorting (HOPS) complex that play a role in the endosome-to-vacuole pathway (24–26). The SM protein Vps45 interacts with and stabilizes the SNARE Tlg2 and acts on the TGN-early endosome pathway (27–30). Vps45 also operates at the late endosome in collaboration with the SNARE Pep12p; this process has also been reported in mammalian cells and in *Aspergillus nidulans* (27,30–32). In this study, we characterized the SM protein homologue Vps45 in *C. neoformans* and discovered roles for the protein in the uptake of exogenous iron, intracellular sorting of an iron uptake protein, mitochondrial function and virulence.

## RESULTS

The *C. neoformans* gene locus CNAG_03628.2 encoding a candidate Vps45 protein was first identified by a reciprocal BLASTp search using the amino acid sequence of *S. cerevisiae* Vps45. The predicted polypeptide (686 aa) from *C. neoformans* displayed 36% identity and 56% similarity to the *S. cerevisiae* protein and 47% identity and 63% similarity to the corresponding protein from *Aspergillus nidulans.* A phylogenetic analysis for Vps45 sequences is presented in Fig. 1. Considering the strong relatedness between orthologs of *VPS45*, we proposed to change the annotation of CNAG_03628.2 to Vps45. To examine the role of Vps45 in *C. neoformans*, we constructed two independent targeted deletion mutants and corresponding strains in which the mutation was complemented with the wild-type (WT) gene. The genotypes of the strains were confirmed by PCR and genome hybridization (S1 Fig.).

**Fig 1.**
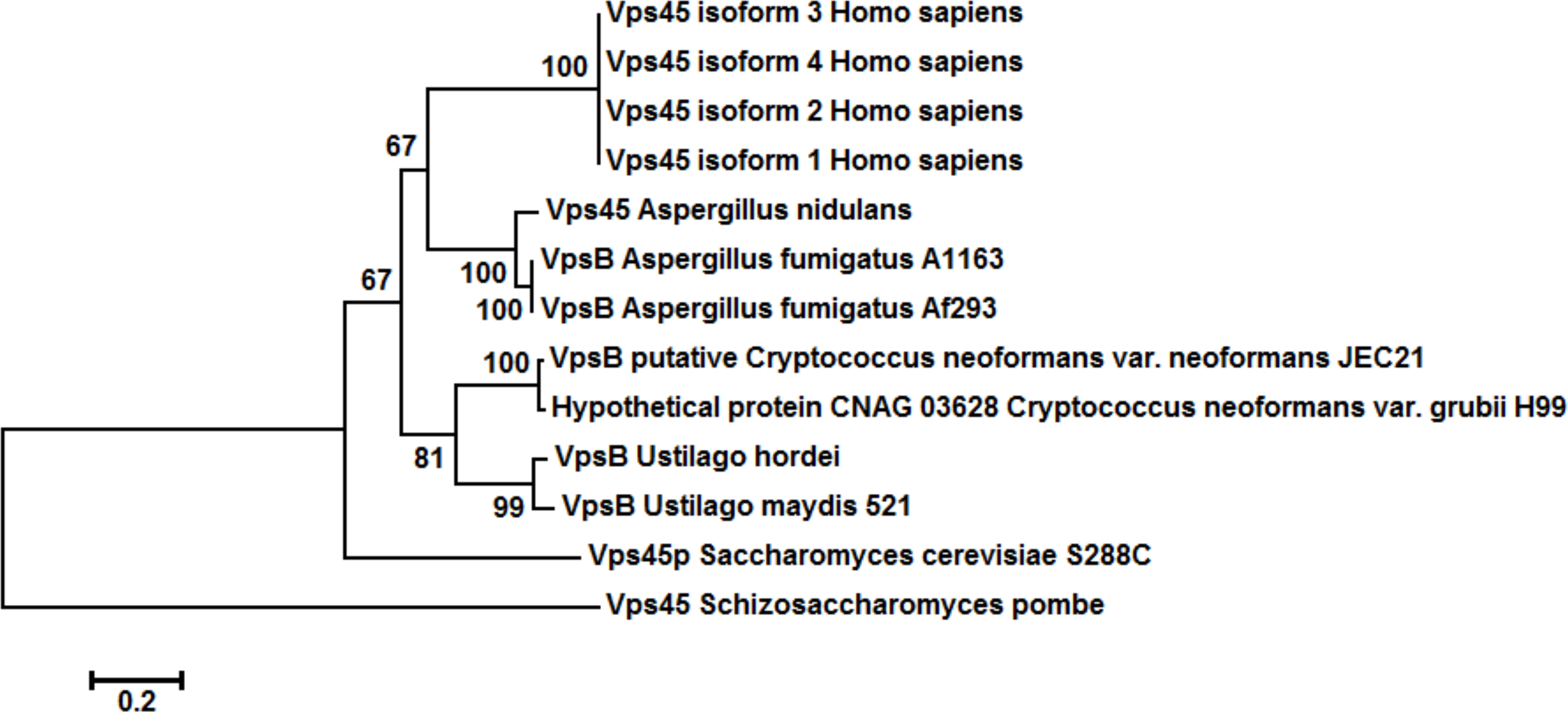
Phylogenetic analysis of the predicted amino acid sequence of Vps45 or VpsB in selected fungi and humans by the Neighbor-Joining method. A phylogenetic tree for Vps45 or VpsB was constructed with orthologous sequences retrieved from NCBI. An unrooted tree was created using the neighbor-joining consensus trees based on the calculated distances using 1,000 bootstrap replications. The evolutionary distances were computed using the Poisson correction method and are in the units of the number of amino acid substitutions per site.

### Vps45 is required for membrane trafficking to the vacuole

As mentioned above, endomembrane trafficking is conducted by specific membrane fusion events that occur via the formation of SNARE bundles, with regulation by SM proteins. To determine whether Vps45 plays a role in endocytosis in *C. neoformans*, cells were grown in low iron conditions, transferred to high iron media, stained with the endocytic tracker dye FM4-64, incubated at 30°C and 37°C, and observed for 90 min. At 30°C, accumulation of the dye on endosomes and at the vacuolar membrane was detected in the WT and complemented strains within 15 min. However, this accumulation at the vacuole was delayed in *vps45* mutants by 15 min. An increase of disorganized cytoplasmic membrane material was also observed over time in mutant cells as opposed to a more organized formation of endosomes observed in WT and complemented strains (Fig. 2A). Similar results were observed when the assay was performed at 37°C, with a general delay in vacuolar staining. Endosomes were observed in WT and complemented strains only after 45 to 90 min while none were detected in mutant strains (S2A Fig.). Moreover, endocytosis assays with FM4-64 were also performed using the *VPS45*-GFP complemented mutant grown at 30°C and 37°C (S2B Fig.). At 30°C, Vps45-GFP fusion protein was first localized at the plasma membrane at 0 min and then to the vacuolar membrane within 15 min (Fig. 2B). Vacuolar colocalization of Vps45-GFP was found after 30 min of incubation at 37°C (S2B Fig.). Notably, introduction of the *VPS4*-5GFP gene fusion into the *vps45* mutant complemented growth defects for a number of conditions at 30°C and 37°C (S3 Fig.).

**Fig 2.**
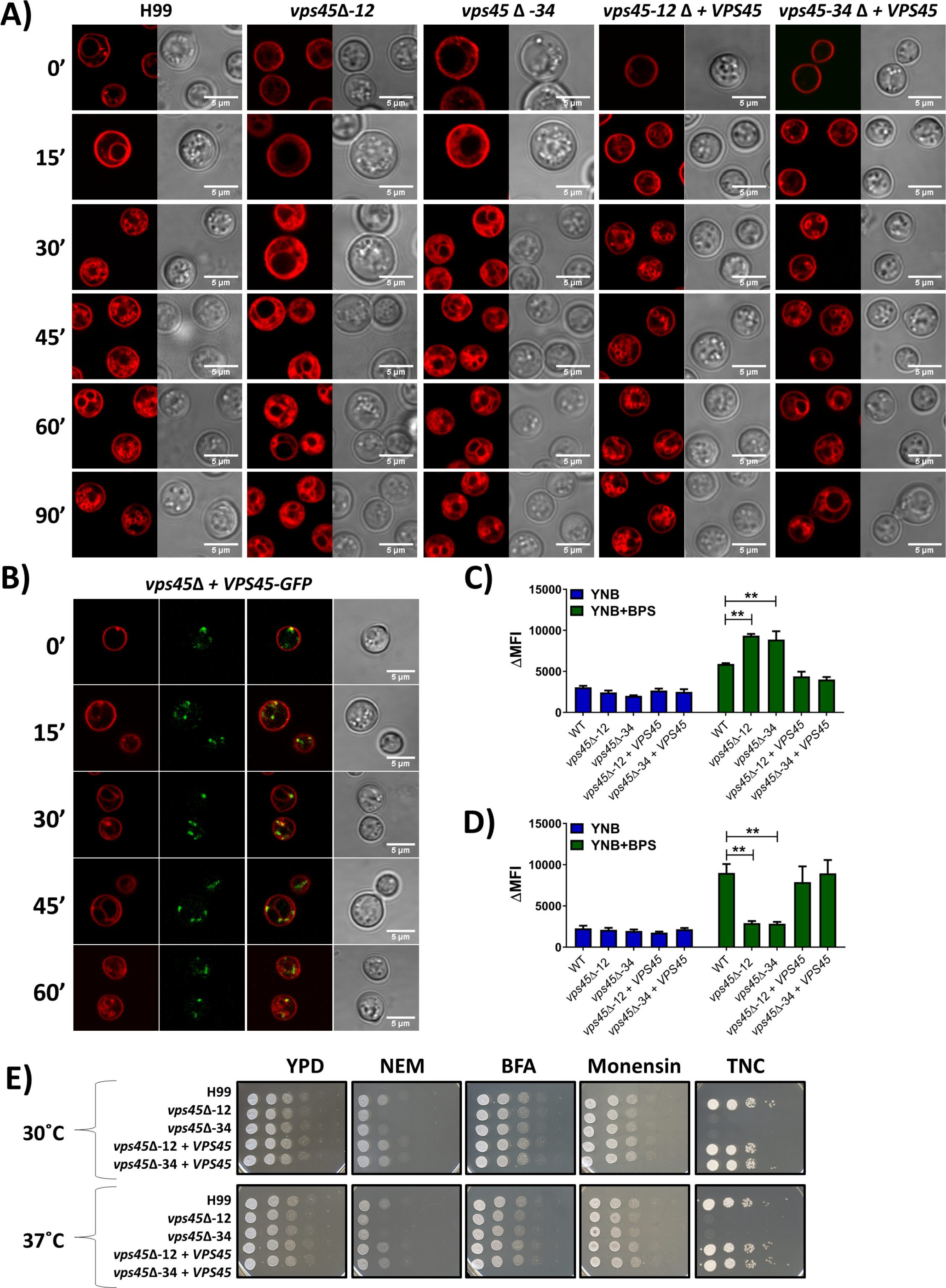
Analysis of endocytosis by FM4-64 internalization. A, B) Uptake of FM4-64 to assess endocytosis in WT, *vps45* mutant, complemented (A), and GFP-tagged (B) strains grown for 24h in YNB-BPS at 30°C. Briefly, 1 × 10^6^ cells/mL were inoculated in YNB-BPS + 100μM FeCl3, stained with 5 μM FM4-64 and transferred in a chamber slide where the cells were maintained at 30°C. Confocal microscopy images were taken every 15 min for 90 min. Colocalization analyses using the ImageJ coloc2 test revealed a positive correlation between VPS45-GFP and endocytic membranes as determined by Pearson’s R value (0.27 - 0.40) and Cortes P-value (0.99-1.00). Intracellular acidification was assessed by flow cytometry on WT, *vps45* mutants and complemented strains grown in YNB and YNB+150μM BPS at 30°C (C) and 37°C (D) and stained with carboxy-DCFDA. The differences in mean fluorescence intensity (ΔMFI) are shown. E) Growth of the WT, mutant and complemented strains was assessed in the presence of trafficking and glycosylation inhibitors. Cells were pre-cultured in YPD overnight at 30°C, serial diluted, and 5μL were spotted onto YPD plates containing 500nM *N*-Ethylmaleimide(NEM), 20μg mL^−1^ brefeldin A (BFA), 500μg mL^−1^ monensin, or 150ng mL^−1^ tunicamycin (TNC). Plates were incubated at 30°C or 37°C for 2 days.

Intracellular acidification due to endosome trafficking is the hallmark of endocytosis (14). Intracellular acidification was assessed by flow cytometry using the acidic pH sensor carboxy-DCFDA dye. WT, mutants and complemented cells did not display any fluorescence differences when grown in YNB at 30°C and 37°C. However, when grown in iron chelated-conditions, *vps45* mutants display increased and reduced fluorescence at 30°C and 37°C, respectively, in comparison to WT and complemented strains (Fig. 2C and D). This suggests that an increase in intracellular pH occurs under iron-depleted conditions at 30°C in *vps45* mutants compared to WT and the complemented strains. However, an increase of intracellular pH was only observed in WT and complemented strains when grown in iron-chelated conditions at 37°C. These results indicate that endocytic processes mediated by Vps45 operate differently depending on the temperature, a finding that is also consistent with the reduce rate of endocytosis and endosome formation previously observed at 37°C. In addition, the mutants, WT and complemented strains were tested for their susceptibility to trafficking inhibitors (i.e., *N*#-Ethylmaleimide (NEM), brefeldin A (BFA) and monensin) and the glycosylation inhibitor tunicamycin (TNC) since many secreted or membrane-bound proteins are *N*-glycosylated. Brefeldin A and monensin did not impair the growth of any of the strains, but the *vps45* mutants showed increased sensitivity to the trafficking inhibitor NEM and the glycosylation inhibitor tunicamycin (Fig. 2E). Taken together, these results are consistent with a role for Vps45 in endocytic events.

### Vps45 contributes to iron acquisition in a temperature dependent manner

We have previously demonstrated that iron acquisition from exogenous heme requires the ESCRT machinery, and this observation implies that intracellular transport of heme occurs by endocytosis (12,13). ESCRT complexes recognize ubiquitinated cargo proteins at endocytic vesicle membranes and are sequentially recruited to deliver endosomes containing extracellular material to the vacuole (33). Therefore, we investigated the role of Vps45 in the ability of *C. neoformans* to use different iron sources. The WT strain, deletion mutants, and the complemented strains were examined for growth in iron-chelated medium (YNB-BPS) (Fig. 3A). After 48h of iron starvation in a pre-culture at 30°C, all strains were subsequently unable to grow in YNB-BPS, but robust growth in YNB with iron was observed, indicating that iron starvation did not kill the cells (Fig. 3A). Furthermore, all of the strains grew at 30°C in YNB-BPS supplemented with 10 μM and 100 μM FeCl_3_ (Fig. 3A). However, when incubated at 37°C, the *vps45* mutants exhibited a delay in growth on YNB-BPS with 25 μM or 100 μM FeCl_3_ (Fig. 3B). This result indicates that Vps45 is required for the use of FeCl_3_ at the host temperature (37°C), but not at lower temperature. We speculate that another system may contribute to iron uptake at 30°C but may be insufficient at 37°C. Similar results were obtained with spot assays on solid media (Fig. 3C). However, when incubated with organic iron sources (i.e. heme, hemoglobin, transferrin and sheep blood), the *vps45* mutants exhibited a delay in growth at both 30°C and 37°C, as compared with WT and complemented strains (Fig. 3A and B). Also, growth was inhibited for the *vps45* mutants when incubated with hemoglobin and blood at 37°C. Again, similar results were obtained with spot assays on solid media (Fig. 3C). These results suggest that Vps45 is required for the use of organic iron sources found in mammalian hosts. Interestingly, when intracellular iron content was measured by inductively coupled plasma mass spectrometry (ICP-MS) for cells grown in high iron conditions at 37°C, the *vps45* mutants displayed a greater amount of iron compared to the WT and complemented strains (Fig. 3D). We hypothesize that the ability to properly traffic the components of iron uptake systems may be required at 37°C. That is, loss of Vps45 may impede the delivery of iron to the vacuole (and sensing of iron repletion) to create an imbalance in iron homeostasis leading to enhanced iron uptake and the increase in intracellular iron content in *vps45* mutants. Altogether, the results indicate that Vps45 is required for intracellular iron transport in a temperature-dependent manner.

**Fig 3.**
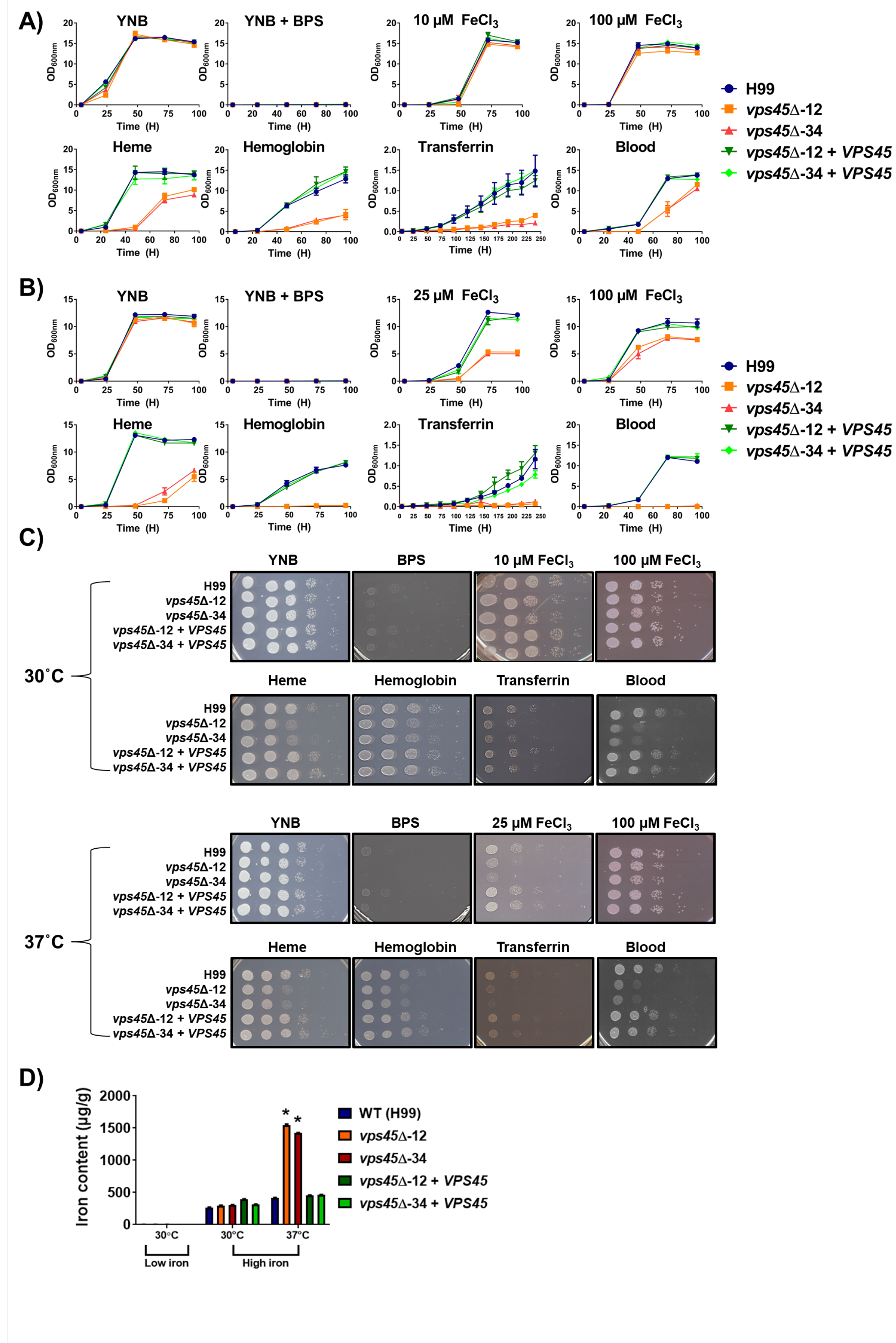
Growth of the WT, *vps45* mutants and complemented strains in inorganic and host-related iron sources. A) and B) Growth assays in liquid media with strains pregrown in YNB+150μM BPS for 48h. After pre-growth, 1×10^5^ of iron-starved cells/mL were inoculated in iron limited conditions supplemented with inorganic iron (i.e. YNB, YNB + 150μM BPS, YNB-BPS + 10μM FeCls (30°C) and + 25μM FeCfe (37°C), YNB-BPS + 100μM FeCl_3_) and organic iron (i.e. YNB-BPS + 10μM Heme, + 2μg mL^−1^ Hemoglobin, + 50μg mL^−1^ Transferrin, + 0.05 % sheep blood). The optical density (600nm) was recorded over a period of 5 days. C) Spot assays are shown for serial diluted WT, mutants and complemented strains in the same conditions as in liquid assays. Plates were incubated at 30°C and 37°C for 2-10 days. D) Inductively coupled plasma mass spectrometry (ICP-MS) measurements of iron content of pre-grown iron starved cells inoculated for 48h in YNB + 150μM BPS w/o 100μM FeCl_3_ at 30°C and 37°C.

### Vps45 contributes to endocytic transport of the ferroxidase Cfo1 for high-affinity iron uptake

In *C. neoformans,* iron from FeCl3 or transferrin is transported by the high-affinity iron uptake system composed of the iron permease Cft1 and the ferroxidase Cfo1 (9,10). Indeed, deletion of either *CFT1* or *CFO1* results in severe growth defects with FeCl_3_ or transferrin as sole iron sources, but not with heme or hemoglobin, which suggests that there are distinct uptake pathways for these iron sources (9,10). The Cft1-Cfo1 system is homologous to the iron permease/ferroxidase Ftr1-Fet3 in *S. cerevisiae,* which was demonstrated to be endocytosed in the presence of iron (19). Given that the Cfo1-GFP fusion protein was found at the plasma membrane and that the *vps45* mutants showed a growth defect in FeCl_3_, we next tested whether the trafficking of Cfo1-GFP was impaired in absence of *VPS45*. As shown in Figure 4 and S4 Fig., Cfo1-GFP started to accumulate at the plasma membrane within an hour of incubation at 30°C (Fig. 4A) and 37°C (S4A Fig.) when cells were transferred from a rich medium (YPD) to the iron-chelated medium (YNB-BPS). These results are consistent with our previous analysis of the location of the Cfo1-GFP protein in the background of a *cfo1*Δ mutant (9). Accumulation of Cfo1-GFP at the plasma membrane was not affected by deletion of *VPS45* suggesting that Vps45 does not play a role in the transport of *de novo* synthesized Cfo1 to the plasma membrane. Furthermore, Cfo1-GFP was located in vesicular bodies that resemble endosomes in the WT and complemented strains, but not in the *vps45* mutant. After 2h of incubation, Cfo1-GFP started to accumulate inside the vacuole of the WT and complemented strains at 30°C (Fig. 4A). Vacuolar localization was maintained after 24h of incubation in low iron media (Fig. 4B). In contrast, Cfo1-GFP was retained at the plasma membrane and dispersed in the cytoplasm in the absence of Vps45, but was never located at the vacuole at the time (3h) and conditions (30°C and 37°C) tested (Fig. 4 and S4A Fig.).

**Fig 4.**
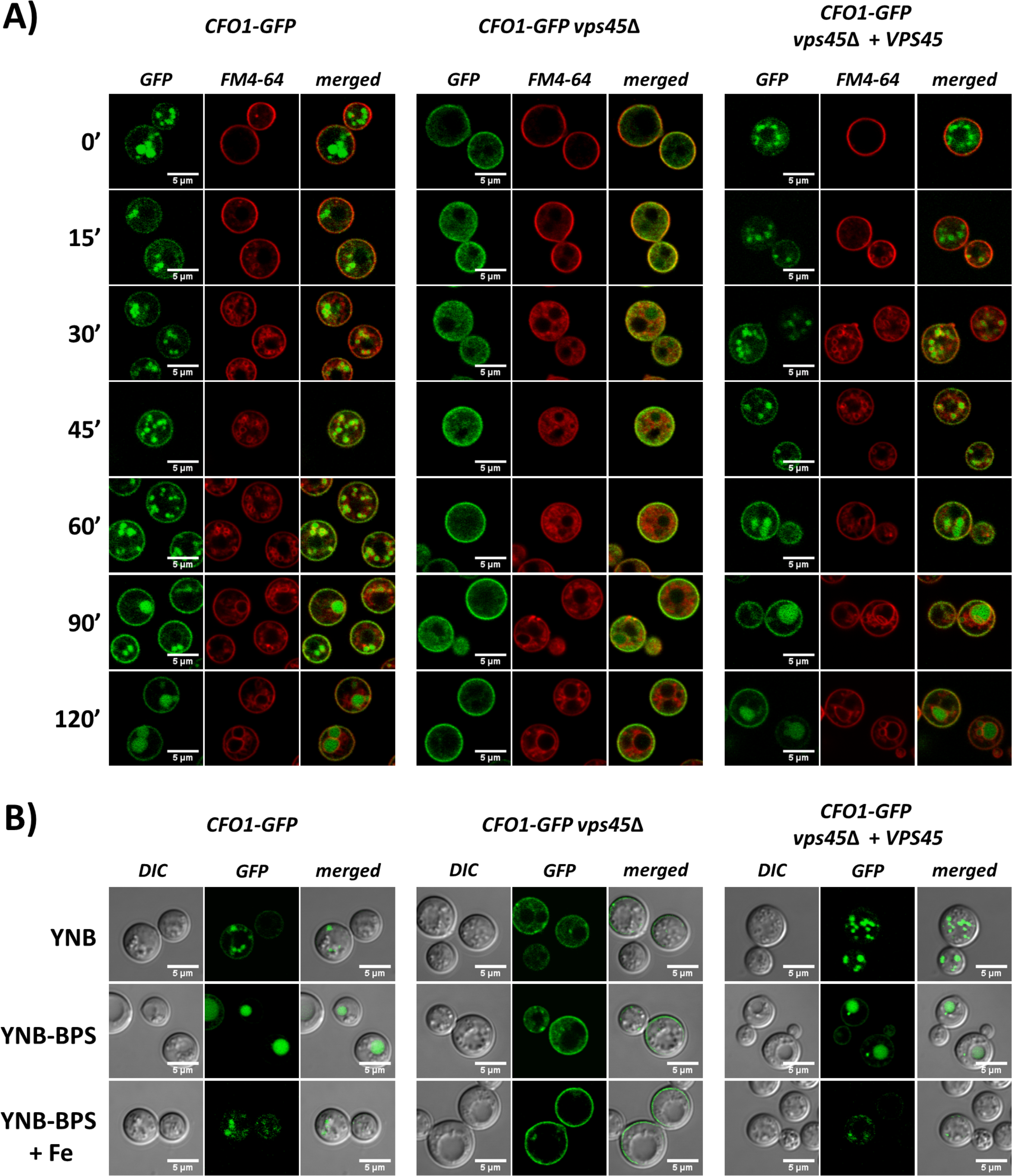
Localization of Cfo1-GFP in the presence and absence of Vps45. A) Localization of Cfo1-GFP in low iron medium. Strains (WT, *vps45*Δ and complement) containing a *CFO1-GFP* construct were cultured overnight in YPD, washed 3 times and counted. Subsequently, 1×10^6^ cells/mL were inoculated in YNB-BPS and incubated at 30°C for 1h. Then cells were stained with 5 μM FM4-64, transferred in a chamber slide and maintained at 30°C. Confocal images were taken every 15 min for 2h. B) Localization of Cfol-GFP after 24h incubation in YNB, YNB + 150μM BPS, and YNB-BPS + 100μM FeCl_3_ at 30°C. The GFP label indicates evaluation of the green fluorescent protein and DIC indicates differential interference contrast microscopy.

Upon addition of FeCl_3_, the Cfo1-GFP signal was reduced in the WT and complemented strains, and was found mostly in vesicular bodies after 24h of incubation at 30°C (Fig. 4B). The signal was also found in vesicular bodies in cells from the YNB medium at 30°C and, as mentioned above, the signal was mainly in the vacuole in cells from YNB-BPS (iron-chelated) medium. With incubation at 37°C in YNB-BPS+FeCl_3_, a strong signal was detected for Cfo1-GFP, and the fusion protein was found at the plasma membrane, in punctate structures and at the vacuole in the WT and complemented strains (S4B Fig.). In these strains, the signal was also found in the vacuole and vesicular bodies in cells from the YNB medium at 37°C, and the signal was mainly in the vacuole in cells from YNB-BPS medium. Again, Cfo1-GFP was located mainly at the plasma membrane in *vps45* mutants when grown in YNB, YNB+BPS and YNB-BPS+FeCl_3_ at 37°C for 24h (S4B Fig.). We established that full length Cfo1-GFP protein was present in the WT, mutant and complemented cells incubated for 3h in YNB-BPS at 30°C and 37°C, as visualized by Western blot (S4C, D Fig.). Taken together, our findings suggest that the intracellular trafficking and expression levels of Cfo1 may be regulated by iron availability and temperature, and that Vps45 plays a role in intracellular transport of the protein from the plasma membrane to the vacuole.

### Vps45 co-localizes with mitochondria, and is required for mitochondrial function and resistance to reactive oxygen species

A *vps45* mutant in *S. cerevisiae* shows increased sensitivity to the antiarrhythmic drug aminodarone that mediates disruption of calcium homeostasis (34). Additionally, mitochondria play key roles in metabolism including iron and calcium homeostasis, and interact with other organelles such as the ER to carry out their functions through communication via diffusion, vesicular transport and direct contact (35,36). These findings, and the involvement of Vps45 in iron homeostasis at least partially through uptake and through endocytosis of Cfo1, prompted us to investigate the potential participation of Vps45 in mitochondrial function. We first determined whether Vps45 associates with mitochondria by examining the strain expressing Vps45-GFP after growth in YNB under iron-depleted conditions at 30°C and 37°C. Mitochondrial staining with mitotracker revealed that the Vps45-GFP protein was co-localized with a subset of mitochondria under every condition tested (Fig. 5A, B). This association occurred in 47% of cells grown in YNB and 58% in YNB+BPS at 30°C. Moreover, temperature elevation increased this association. Indeed, 58% of cells grown in YNB and 75% of cells grown in YNB-BPS at 37°C were found to show Vps45 in association with mitochondria. Moreover, observations of mitochondria morphology did not revealed any differences between WT, mutants and complemented strains under iron-depleted conditions at 30°C and 37°C (S5A, B Fig.). In parallel, the WT, mutant and complemented strains were tested for their susceptibility to inhibitors of the mitochondrial electron transport chain (ETC) complexes. Growth was not impaired for any strains in the presence of inhibitors of ETC complexes I and II (i.e., rotenone, malonic acid, and oligomycin A) (S5C Fig.). However, the *vps45* mutants showed increased sensitivity to the alternative oxidase inhibitor salicylhydroxamic acid (SHAM), the complex III inhibitor antimycin A, and the complex IV inhibitor potassium cyanide (KCN), when compared to the WT and complemented strains (Fig. 5C). Increased sensitivity to hydrogen peroxide (H2O2) was also observed for the *vps45* mutants, but no growth defect was noted when the mutants were exposed to other agents that caused oxidative stress such as paraquat, plumbagin, diphenyleneiodonium chloride (dpi) and menadione (Fig. 5C, and S5C Fig.).

**Fig 5.**
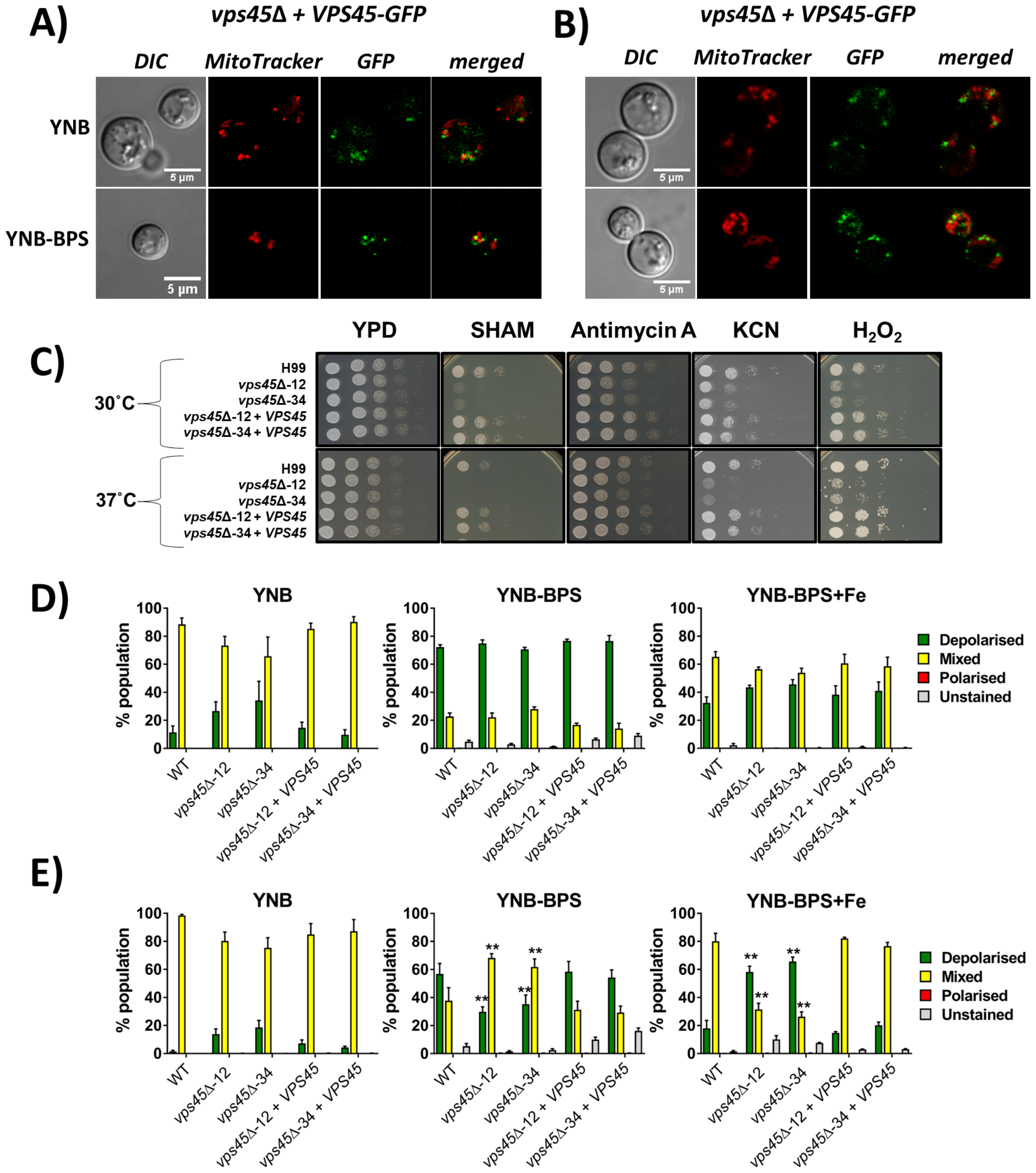
Impact of Vps45 on mitochondrial function. A) and B) The association of Vps45-GFP with mitochondria was examined with the *vps45* mutant complemented with *VPS45-GFP.* The strain was cultured in YNB and YNB + 150μM BPS for 24h at 30°C (A) and 37°C (B), cells were stained with 500nM MitoTracker Red CMXRos (mitotracker) and images were taken with a confocal microscope. C) The involvement of Vps45 in mitochondrial function and ROS sensitivity was examined by spot assays on inhibitors and H_2_O_2_. The WT, mutants and complemented strains were pre-cultured in YPD overnight at 30°C, serial diluted, and 5μL were spotted onto YPD plates containing either 5mM salicylhydroxamic acid (SHAM), 3μg mL-1 antimycin A, 10mM potassium cyanide (KCN) or 0.01% H_2_O_2_. Plates were incubated at 30°C and 37°C for 2 days. D) Mitochondrial membrane potential was assessed by flow cytometry on 50,000 cells stained with JC-1 (5μM) and grown in iron deplete and replete conditions at 30°C (D) and 37°C (E). Experiments were carried out at least two times in triplicate (ANOVA ** *P* < 0.001).

Mitochondrial membrane potential (MMP) was also measured by flow cytometry in cells stained with JC-1. The JC-1 dye exhibits potential-dependent accumulation in mitochondria as indicated by a fluorescence emission shift from green (~ 529 nm) to red (~ 590 nm). Consequently, mitochondrial depolarization is indicated by a shift from red to green fluorescence (S5D Fig.). The WT, mutant and complemented strains were grown in YNB under iron-depleted and replete conditions at 30°C and 37°C, and stained with JC-1 for 30 min. Mitochondria membrane potential was assessed for each strain by determining the % of the cells in the population with polarized (only red signal), depolarized (only green signal) and mixed polarization (red and green signal). When grown in YNB at both 30°C and 37°C, all cells displayed a mixture of polarized and depolarized mitochondria membrane potential. However, upon iron depletion at 30°C, a shift from mixed polarization to a fully depolarization population was observed and restoration of the mixed polarization population was noted under iron-replete conditions (Fig. 5D). Conversely, under the iron-depleted condition at 37°C, *vps45* mutants maintained a mixture of polarized and depolarized mitochondrial membrane potential while WT and complemented strains showed an increase in the depolarized population. Addition of iron restored mixed polarization in the WT and complemented strains, while a shift towards depolarization in *vps45* mutants was detected (Fig. 5E). These results suggest that iron homeostasis impacted mitochondria membrane potential in the *vps45* mutants at 37°C. Since iron uptake, trafficking and sensing are impaired in the absence of Vps45, especially at high temperature where cells displayed a greater amount of intracellular iron, we speculate that iron accumulation in these mutants contributes to their mitochondria membrane depolarization. Paradoxically, chelation of iron by the addition of BPS may alleviate the potential dysfunction in iron homeostasis, and this may translate into maintenance of mitochondrial membrane potential.

### Vps45 is involved in calcium homeostasis

Along with the ER, mitochondria also play important roles in calcium homeostasis and signaling (37). Given the connections between Vps45 and mitochondrial function established above and known connections with calcium homeostasis in yeast (34), we next tested our strains for their susceptibility to calcium inhibitors. Growth was not impaired for the WT and complemented strains in the presence of aminodarone, cyclosporine A, tacrolimus (FK506), and the extracellular calcium chelator EGTA, but the *vps45* mutants showed increased sensitivity to these compounds (Fig. 6A). Moreover, chelation of calcium with EGTA resulted in growth inhibition for all strains in liquid media (Fig. 6B). Addition of calcium restored the growth of the strains when incubated at 30°C, but the growth of the *vps45* mutants remained hindered at 37°C (Fig. 6B). This result suggests that Vps45 is required for calcium use at host temperature. Mitochondrial morphology was assessed in cells grown in calcium-chelated conditions at 30°C and 37°C (Fig. 6C and D). A mixture of puncta and rod shaped mitochondria were observed when cells were grown in YNB at 30°C or 37°C, and no marked change in mitochondrial morphology was observed when cells were exposed to the calcium chelator EGTA at either temperature.

In light of our finding that Vps45 participates in mitochondrial functions, we next determined whether Vps45 associated with mitochondria under calcium-limited conditions. The strain expressing the *VPS45-GFP* construct was grown in YNB under calcium-depleted conditions at 30°C or 37°C. Mitochondrial staining with mitotracker showed that Vps45-GFP protein was found associated with a subset of mitochondria upon calcium restriction in 61% of cells grown at 30°C and 87% of cells grown at 37°C (Fig. 6E and F). We also measured the mitochondrial membrane potential (MMP) by flow cytometry in cells stained with JC1-1 (S5D Fig.). When the WT and complemented strains were grown under calcium-starved conditions (YNB-EGTA) at 30°C, a shift from mixed polarized to depolarized mitochondria was observed. A similar shift was also obtained for *vps45* mutants but to a lesser extent. Interestingly an increased in temperature restored mitochondrial membrane potential population to levels similar to those found when cells were grown in YNB at 37°C (Fig. 6G). We went on to measure intracellular calcium content by inductively coupled plasma mass spectrometry (ICP-MS), and found that only the *vps45* mutants displayed a greater amount of calcium compared to WT and complemented strains in high iron conditions at 37°C (Fig. 6H). Taken together, these results suggest that Vps45 influences calcium homeostasis and mitochondrial membrane potential in a manner related to iron availability.

**Fig 6.**
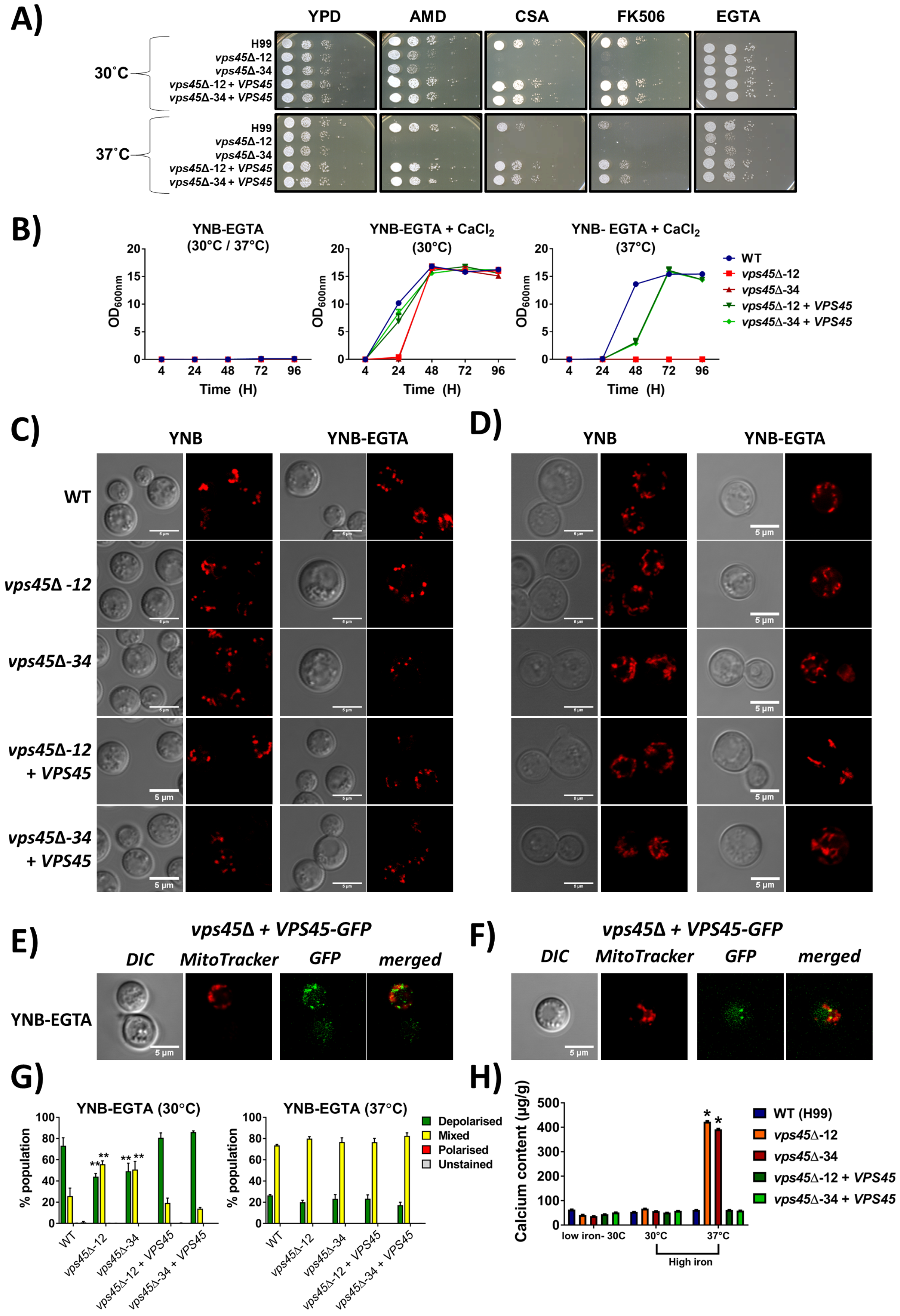
Vps45 is involved in calcium homeostasis. A) To examine the calcium-related phenotypes for the *vps45* mutants, the WT, mutants and complemented strains were precultured in YPD overnight at 30°C, serial diluted, and 5μL were spotted onto YPD plates containing either 25μM aminodarone, 100μg mL-1 cyclosporine A (CSA), 500ng mL-1 tacrolimus (FK506), or 5mM EGTA. Plates were incubated at 30°C and 37°C for 2 days. B) Growth curves of the WT, mutant and complemented strains inoculated in calcium-chelated media. Strains were pre-grown in YNB-EGTA for 48h and 1×10^5^ of calcium-starved cells/mL were inoculated in YNB-EGTA, YNB-EGTA + 50mM calcium and incubated at (30°C) and (37°C). Optical density (600nm) was recorded over a period of 5 days. Disruption of *VPS45* influences mitochondria morphology in calcium chelated conditions. Confocal images of mitochondria stained with 500nM mitotracker were taken of the WT, mutant and complemented strains inoculated for 24h in YNB ± 50mM EGTA at 30°C (C) and 37°C (D). Vps45 associates with mitochondria in calcium chelated conditions. The *vps45* mutant complemented with *VPS45-GFP* was cultured in YNB + 50mM EGTA, for 24h at 30°C (E) and 37°C (F) and stained with mitotracker. Mitochondrial membrane potential was assessed by flow cytometry on 50,000 cells stained with JC-1 (5μM) and grown in calcium deplete conditions at 30°C and 37°C (G). Experiments were carried out at least two times in triplicate (ANOVA ** *P* < 0.001). H) Inductively coupled plasma mass spectrometry (ICP-MS) measurements of the calcium content of pre-grown iron starved cells inoculated for 48h in YNB + 150 μM BPS w/o 100μM FeCl_3_ at 30°C or 37°C. The same cells were used for iron measurements by ICP-MS in figure 3D.

### Vps45 influences cell wall integrity and capsule formation in a temperature-dependent manner

Our analysis of the role of Vps45 in iron and calcium homeostasis indicated that mutant phenotypes were generally exacerbated at elevated temperature. Given that the transcription factor Crz1 is an effector of calcineurin, a key mediator of calcium signaling that is activated by temperature stress (38), we further examined phenotypes related to calcineurin and Crz1 function. In particular, genes involved in cell wall remodelling are regulated by Crz1 in *C. neoformans* and we therefore investigated whether *vps45* mutants were sensitive to agents that challenge cell wall integrity (38,39). Specifically, we exposed our strains to calcofluor white, SDS, caffeine or NaCl, and found that the *vps45* mutants displayed growth defects, especially at 37°C, when compared to the WT and complemented strains (Fig.7A). Furthermore, deletion of *CRZ1* in a *vps45* mutant abolished growth on calcofluor white and caffeine at 37°C (Fig. 7B). In addition, the double knockout mutants displayed a growth defect on YPD at 37°C, and when grown on the calcium chelator EGTA or the drug aminodarone that disrupts calcium homeostasis. These results suggest a role for Vps45 in calcium signaling that may overlap with functions controlled by the calcineurin and Crz1 pathway. To further investigate cell wall remodelling in the absence of *VPS45*, we stained the cells with the fluorescent probe pontamine, a non-specific cell wall dye, and with lectin-binding probes, such as eosin Y and concanavalin A, which respectively bind chitosan and mannans present in the cell wall. By measuring differential fluorescence intensity by flow cytometry, we found that *vps45* mutants showed increased fluorescence when stained with pontamine and concanavalin A, but not with eosin Y (Fig. 7C). The observed differences were greater when the mutant cells were grown at 37°C and compared to the WT and complemented strains. These observations suggest that defects in cell wall structure result from the absence of *VPS45.*

**Fig 7.**
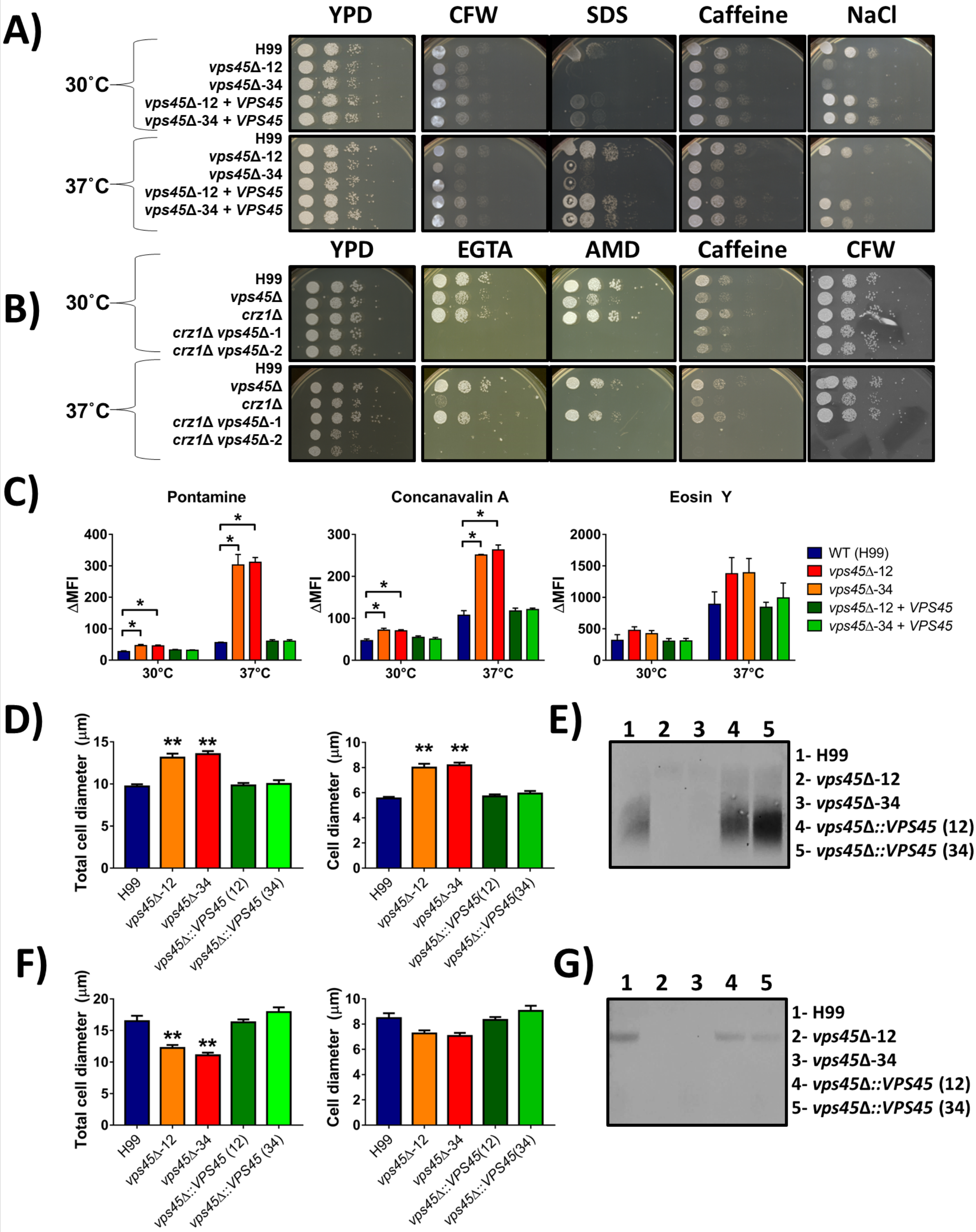
Cell wall integrity is altered in *vps45* mutants. A) Spot assays of serial diluted WT, *vps45* and *crz1* mutants and *vps45* complemented strains grown in the presence of cell wall stressors. Cells were pre-cultured in YPD overnight at 30°C, serial diluted, and 5μL were spotted onto YPD plates containing either 1.0 mg mL^−1^ calcofluor white (CFW), 0.01% sodium dodecyl sulfate (SDS), 0.5 mg mL^−1^ caffeine or 1.5M sodium chloride (NaCl). Plates were incubated at 30°C or 37°C and photographed after 48h of incubation. C) Flow cytometry measurement of differential fluorescence of cells stained with 100μg mL^−1^ pontamine, 100μg mL^−1^ concanavalin A, or 250μg mL^−1^ Eosin Y. Cells were grown in YPD at 30°C and 37°C for 24h and differences in the mean fluorescence intensity (ΔMFI) were determined between stained and unstained samples. Analyses were performed on the gated WT unstained population. Capsule thickness, cell diameter measurements and GXM detection in supernatants (D-G) of cell grown in defined limited iron media (33) at 30°C (D) and 37°C (F). Cells were grown in LIM at 30°C for 48 h and the capsule thickness and cell diameter of at least 40 cells of each strain were measured. Each bar represents the average with standard deviations. Statistical significance relative to WT is indicated by an asterix (ANOVA ** *P* < 0.0001). Shed polysaccharides were collected from culture supernatants of all strains incubated at 30°C (E) or 37°C (G). The electrophoretic mobility and relative quantities of shed GXM were assessed by immunoblotting with the anti-GXM antibody mAb18b7.

Changes in cell wall integrity may also influence the exposure of surface molecules that are important for virulence, such as capsular polysaccharides. We therefore analyzed the impact of the loss of *VPS45* on capsule elaboration (Fig. 7D-G). First, we assessed capsule size by India ink staining of cells cultured in defined limited-iron media for 48h at 30°C (D) or 37°C (F). We observed a significant change in capsule size after deletion of *VPS45* and the change was temperature dependent. That is, capsule and cell size increased when cells of the *vps45* mutants were grown at 30°C (Fig. 7D). However, a reduction in capsule size only was observed at 37°C (Fig. 7F). Glucuronoxylomannan (GXM) is the major polysaccharide of the capsule that is both attached to the cell wall and shed into culture medium (40). Given that cell wall remodelling defects in the *vps45* mutants might influence capsule attachment, we determined whether *vps45* mutants shed different amounts of GXM versus WT cells by performing a capsule blot assay (41,42). The cells were incubated in defined low-iron media for 48h at 30°C or 37°C and the relative abundance of shed polysaccharide was analyzed by reactivity with an anti-GXM antibody (mAb18B7). As shown in Fig. 7E, the *vps45* mutants shed very little GXM into the supernatant compared to WT and complemented strains when grown at 30°C. This result suggests that capsular material is fully attached to the cell wall rather than being secreted into the extracellular space, a result consistent with the finding that cells and capsule are larger in *vps45* mutants when grown at 30°C. Conversely, capsular material was barely detected in the supernatants of cultures from *vps45* mutants when incubated at 37°C (Fig. 7G). Given the reduced capsule size of these cells, this result suggests that capsular material is synthesized at a reduced level and/or that a defect in secretion may occur under these conditions. However, further investigation revealed that mutant cells are unable to survive and proliferate under the conditions of minimal media and high temperature for the experiment, and this could explain the reduced capsular material on the cell surface and in the supernatant (S6 Fig.). Overall, these observations revealed that deletion of *VPS45* caused defects in the capsule formation and cell survival under austere conditions.

### Loss of Vps45 causes increased sensitivity to chloroquine and azole drugs

Chloroquine and quinacrine are antimalarial drugs that accumulate inside lysosomes (vacuoles) of the parasite *Plasmodium* and interfere with hemoglobin digestion. Briefly, ingested hemoglobin is deposited inside the digestive vacuole where degradation occurs and heme is transformed into an inert crystalline polymer called hemozoin. Chloroquine and derivatives accumulate in the vacuole and bind to hematin, a toxic product of hemoglobin degradation, preventing its incorporation into hemozoin, thus intoxicating the parasite (43). On the other hand, azoles, including imidazoles (e.g. miconazole) and triazoles (e.g. fluconazole) interfere with ergosterol biosynthesis by inhibiting the enzyme lanosterol 14-α-demethylase. Because ergosterol is a major constituent of the fungal membranes, its depletion results in growth inhibition (44). Since Vps45 was found to be associated with plasma and vacuolar membranes, and a mutant displayed defects in cell wall integrity, we tested the sensitivity of *vps45* mutants to cell wall, antimalarial, and azole drugs. As shown in Figure 8A, *vps45* mutants display an increased sensitivity to caspofungin, quinacrine, chloroquine and fluconazole when compared to the WT and complemented strains, whereas all strains were sensitive to miconazole. This marked sensitivity is noticeable at both 30°C and 37°C. The drug sensitivity of the *vps45* mutant could be explained by the fact that perturbation of membrane trafficking, fusion events and potentially cargo delivery impairs intrinsic resistance mechanisms. Addition of exogenous heme has been demonstrated to reverse sensitivity to fluconazole in a *cfo1*mutant (9). Therefore, we tested if addition of iron and heme had an impact on drug susceptibility and we found that supplementation of iron but not hemin improved the growth of *vps45* mutants on fluconazole when incubated at 30°C, but not at 37°C (Fig. 8B). This result is in agreement with the previous results that Vps45 is required for the use of FeCl_3_ at the host temperature (37°C), but not at lower temperature. Furthermore, addition of iron and heme to miconazole aided the growth of WT and complemented strains, but not the *vps45* mutants (Fig. 8B). Overall, these results suggest that Vps45 is required for resistance to drugs that target the cell wall, membranes and the vacuole.

**Fig 8.**
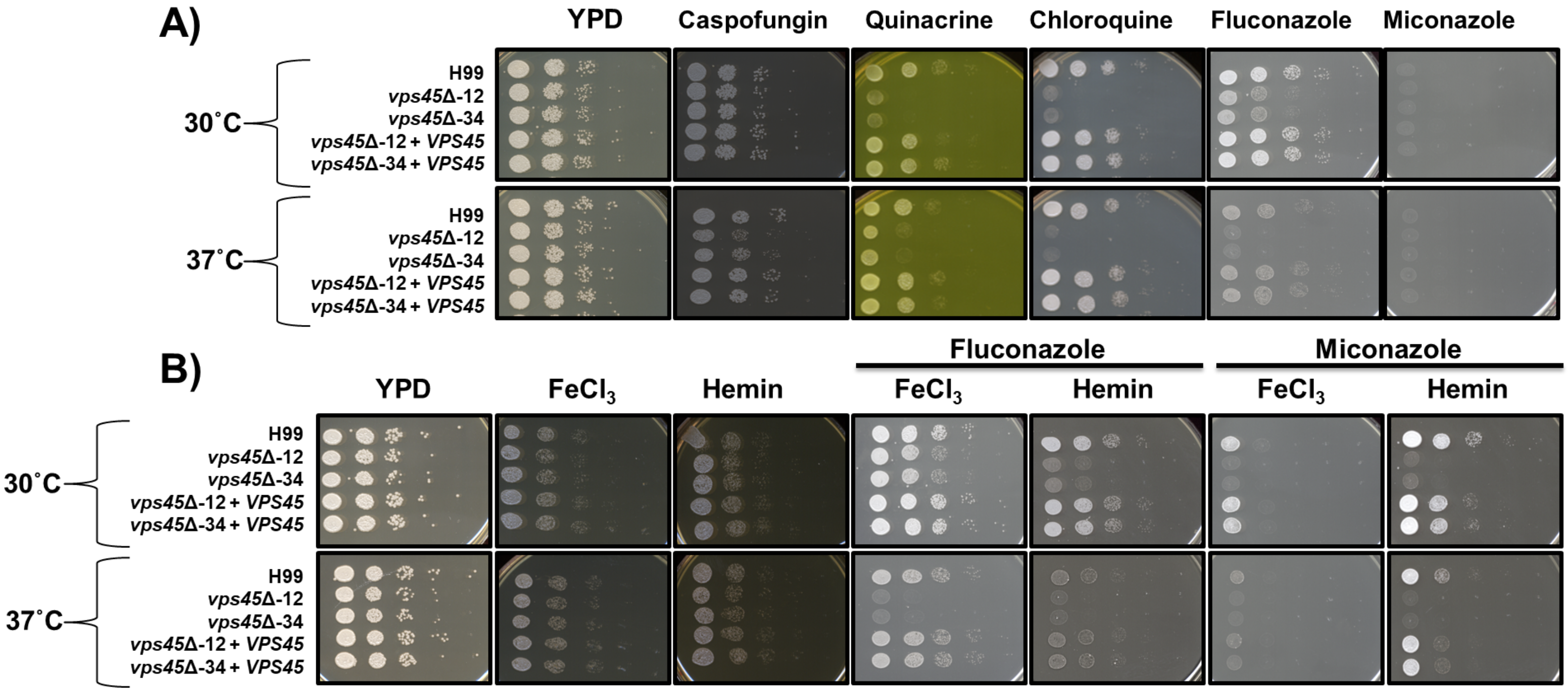
Vps45 is involved in resistance to drugs that target the membrane and vacuole. The WT, mutants and complemented strains were grown in the presence of antimalarial and antifungal drugs. Cells were pre-cultured in YPD overnight at 30°C, serial diluted, and 5μL were spotted onto YPD plates containing either 10 μg mL^−1^ caspofungin, 1.6 mM quinacrine, 6 mM chloroquine, 10 μg mL^−1^ fluconazole and 0.05 μg mL^−1^ miconazole (A). Growth with supplementation of 100 μM feCl_3_ and 100 μM heme to antimalarial and antifungal drugs was assessed (B). Plates were incubated at 30°C or 37°C for 2 days.

### Vps45 is required for virulence in a mouse model of cryptococcosis and for survival in phagocytic cells

Given the contributions of Vps45 to capsule formation and the influence of host temperature on mutant phenotypes, we next tested the importance of *VPS45* for the development of cryptococcosis. We first examined the survival and proliferation of fungal cells upon interaction with murine macrophage-like J774A.1 cells. As shown in Figure 9A, CFU numbers of *vps45* mutants recovered at 24h were significantly lower than for the WT and complemented strains suggesting a reduced ability to survive and proliferate inside macrophages. This result prompted a further assessment of virulence in mice. Groups of 10 female BALB/c mice were inoculated intranasally with WT, a *vps45* mutant or the corresponding complemented strain and monitored daily for disease development. The WT and complemented strains caused a lethal infection in all mice by days 18 and 23, respectively. However, mice inoculated with the *vps45* mutant showed no sign of disease and survived until the end of the experiment (50 days) (Fig. 9B). This avirulent phenotype was confirmed by examination of the fungal loads in organs harvested from the infected mice. As opposed to the mice infected with the WT or complemented strains, no cells of the *vps45* mutant were retrieved from the bloodstream or any of the organs (i.e., lungs, brain, liver, spleen or kidney) (Fig. 9C). Overall, these results indicate that *VPS45* is required for the survival inside macrophages as well as virulence and dissemination to systemic organs in mice.

**Fig 9.**
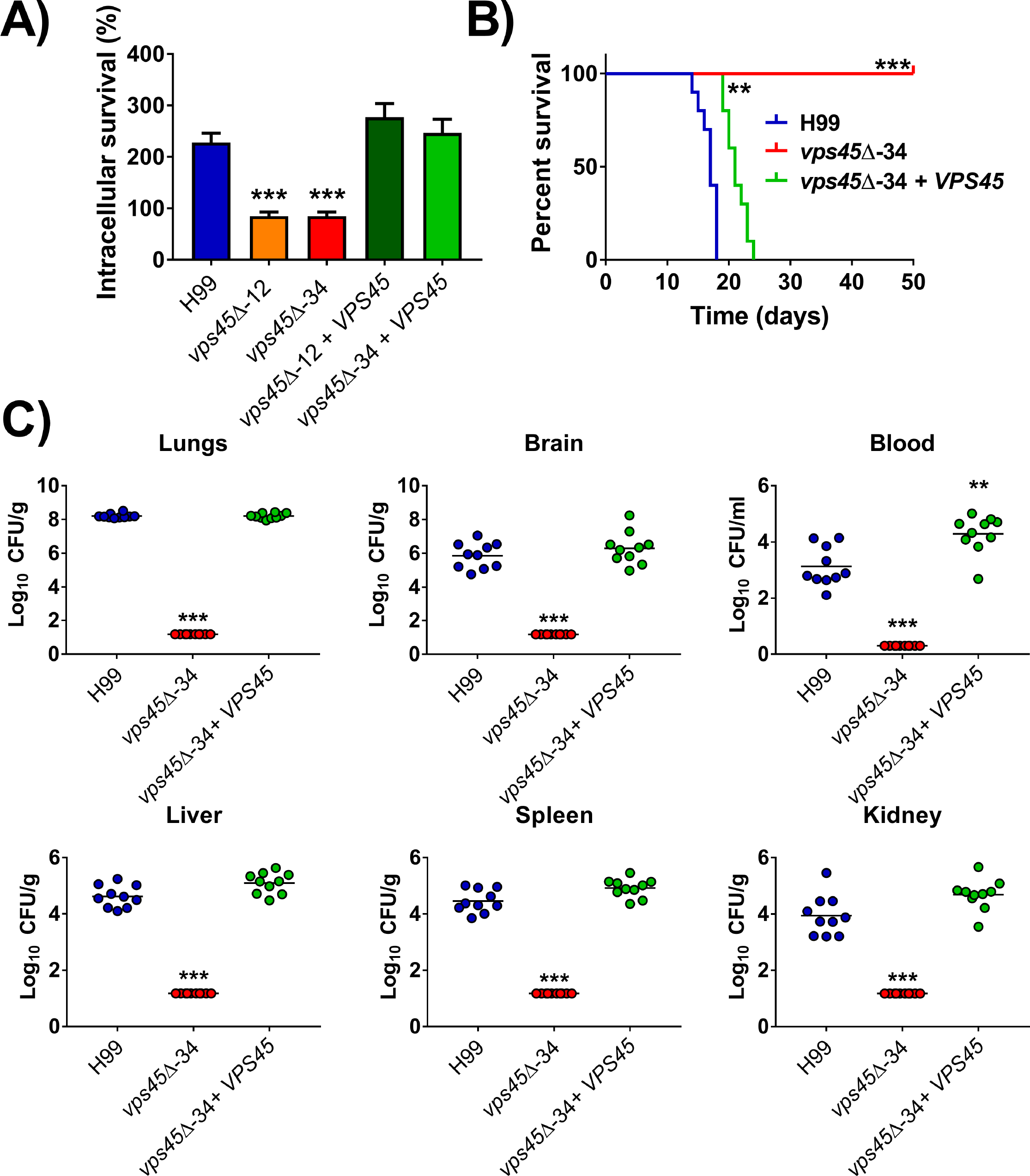
Vps45 is required for survival in macrophages and virulence in mice. A) WT, *vps45* mutants and complemented strains were incubated with the murine macrophagelike cell line J774A.1. The strains were incubated for 2 h and 24 h in DMEM with macrophages at a MOI of 1:1 at 37°C. CFUs from lysed macrophages were obtained and survival was calculated as the ratio of CFUs at 24 h versus 2 h of incubation. The data represent the mean values ± standard error of the mean of four independent biological experiments done in triplicate. Statistical analysis was performed using an unpaired twotailed Student’s t test using WT as the reference strain (*** *P* < 0.0001). B) 10 female BALB/c mice were inoculated intranasally with 2×10^5^ fungal cells and survival of the mice was monitored daily. Survival differences between groups of mice were evaluated by log-rank test against WT. (*** *P* < 0.0001). C) Fungal burden was determined in systemic organs (lungs, brain, liver, kidney and spleen) and in cardiac blood for all mice at time of death. The Mann-Whitney U test was used for statistical analysis using WT as the reference strain (** *P* < 0.005, *** *P* < 0.0001).

## DISCUSSION

The SM protein Vps45 plays an important role in vesicle trafficking by conferring specificity on SNARE proteins that mediate docking and fusion events (27). As initially demonstrated in *S. cerevisiae*, a *vps45* mutant is defective in vacuolar biogenesis due to impaired fusion of Golgi-derived vesicles with the prevacuolar compartment (28,45). In this study, we investigated the role of an ortholog of Vps45 in iron trafficking in *C. neoformans*, and found that the protein is needed for robust growth of the pathogen on both inorganic and organic iron sources. Notably, the iron-related phenotypes were more severe at 37°C. Interestingly, loss of Vps45 changed the intracellular distribution of a fusion protein of GFP with the ferroxidase Cfo1 that mediates high affinity iron uptake in *C. neoformans.* Specifically, the WT strain accumulated Cfo1-GFP at the plasma membrane and vacuole under low iron conditions while the protein was found in the PM and internal punctate structures (presumably endosomes) in a *vps45* mutant. That is, loss of Vps45 prevented vacuolar accumulation. Under high iron conditions, the protein accumulated in internal punctate structures in the WT strain, and in the PM and internal dispersed structures in the mutant. These results suggest that proper trafficking of Cfo1, including localization at the vacuole, contributes to robust growth on FeCl_3_. The differences in localization and growth were most marked at 37°C, and the mutant hyper accumulated iron at this temperature. We speculate that proper trafficking of Cfo1 to the vacuole contributes to robust iron uptake, correct intracellular distribution, and overall homeostasis (Fig. 10).

**Fig 10.**
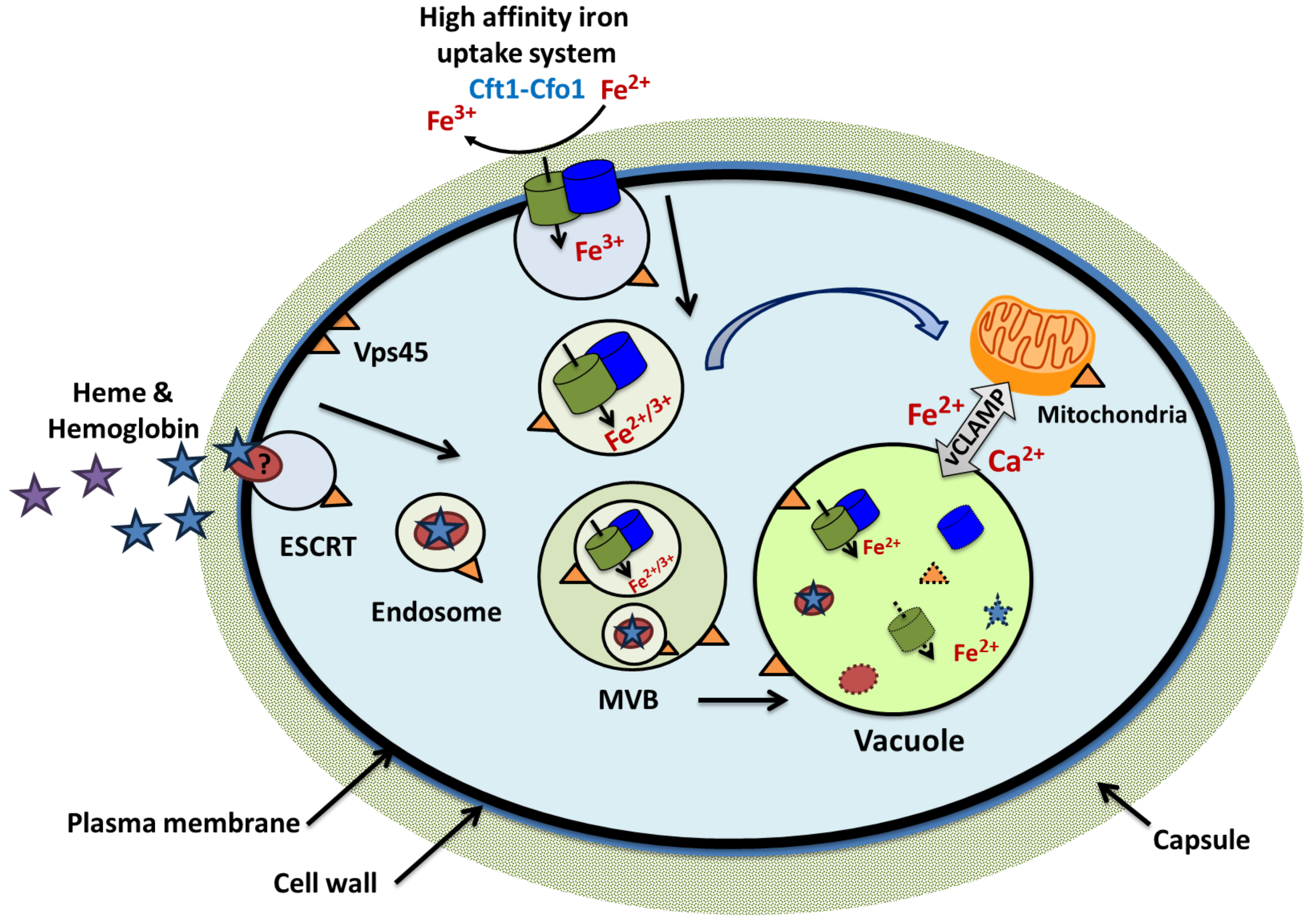
Schematic representation of the role of Vps45 in endocytosis of exogenous iron sources and in intracellular trafficking from the plasma membrane to the vacuole and mitochondria. The internalization of newly created endosomes loaded with the high affinity iron uptake system (Cft1-Cfo1) and heme and a putative receptor (?) is depicted. These endosomes may fuse into a pre-vacuolar compartment also called a multivesicular body (MVB) and undergo progressive acidification of the intraluminal milieu until fusion with the vacuole. Multiple membrane fusion events occur during that process and endosomal cargo is ultimately delivered into the vacuole for dissociation and degradation. Direct fusion from iron-loaded endosome to the mitochondrial membrane is also possible (58), but has yet to be demonstrated in *C. neoformans.* An exchange of iron and calcium ions between the vacuole and mitochondria is likely and may require Vps45 for vCLAMP formation (36).

Our analysis of the Vps45-GFP fusion also revealed an unexpected co-localization with mitochondria, and we subsequently found that *vps45* mutants were sensitive to specific inhibitors of electron transport and to reactive oxygen species. Additionally, the *vps45* mutants appeared to be impaired in calcium homeostasis, and mitochondria are known to be important organelles for calcium storage and signaling. Given that mitochondria-endomembrane system connections are emerging as important features of intracellular communication (36,46), it is tempting to speculate that Vps45 functions in these connections. That is, Vps45 may directly or indirectly contribute to establishing or maintaining membrane contact sites between mitochondria and the vacuole or other endomembranes. Interestingly, contacts contribute to iron and calcium distribution in the cell including trafficking to mitochondria because it has been demonstrated in erythrocytes that iron loaded transferrin is directly transported from the endosomes to mitochondria (47). The observed association of the Vps45-GFP fusion with mitochondria supports a possible direct contribution to membrane contact sites. However, alternative or additional possibilities are that loss of Vps45 might impair the delivery of proteins needed for contact site formation and/or vacuolar function to indirectly influence mitochondrial activities via changes in metabolism, response to oxidative stress and mitophagy (46,48). These ideas are supported by genetic screens in yeast that identified many VPS genes as contributing to mitochondrial morphology (48,49).

The participation of Vps45 in other endosomal trafficking connections and/or impaired vacuolar function may also explain the additional phenotypes of the *vps45* mutant. For example, impaired delivery of proteins and materials to the PM and cell wall may explain increased sensitivity to CFW, Caffeine, SDS and NaCl. Similarly, participation of Vps45 in vesicle transport to the ER or establishment of connections between the ER and the PM, vacuole or mitochondria might explain the increased sensitivities to tunicamycin, chloroquine, quinacrine, the azole antifungal drugs and capsofungin.

The *vps45* mutant was also attenuated for virulence in a mouse model of cryptococcosis and for survival in a macrophage-like cell line. Adaptation to the host environment requires proper nutrient acquisition, and virulence factor elaboration. In this regard, defects in iron acquisition are known to attenuate the virulence of *C. neoformans* (3,9,12). Importantly, the observed cell wall changes and altered capsule elaboration with reduced shedding also likely reduced virulence given the established importance of the capsule in fungal cell protection and immuno-modulatory properties. Capsular polysaccharide shedding (mainly composed of GXM) tends to accumulate in patient serum and cerebrospinal fluid and supress patient immune responses by limiting immune cell infiltration leading to devastating consequences. The release of GXM is regulated by environmental cues such pH and nutrient availability (50). Furthermore many of the phenotypes of the *vps45* mutants are exacerbated at 37°C and this would further preclude robust proliferation in mammalian hosts. Additional phenotypes for signaling (calcium), and mitochondrial function (respiration and ROS maintenance) could also potentially reduce growth in the host as mitochondria metabolism and ROS accumulation are linked to capsule formation and enlargement (51).

Our results support the need for additional work on Vps45 and other SM proteins for *C. neoformans*. In particular, the identification of interacting partners and a more detailed analysis of the location vis-à-vis mitochondria in different growth conditions is needed to more fully understand the contribution of Vps45. Vps45 and Vps33 were recently characterized in the fungus *Aspergillus nidulans* as Sec1/Munc-18 proteins involved in the regulation of the syntaxin PepA^Pep12^ which contributes to membrane compartment identity. The authors demonstrated that Vps45 plays role in regulating _PepA_^Pep12^ in the early endosomes formation whereas Vps33 contributes to the regulation of PepA^Pep12^ in late endosomes/vacuoles (31). Bioinformatics analysis of H99 genome indicated that a homologue of Vps33 does exist (CNAG_02373), but preliminary results revealed no phenotypic defect of a *vps33* mutant for iron uptake (Bakkeren, Hu, Caza, unpublished). However, further investigation is needed to characterize the role of VPS33 in endosome sorting in *C. neoformans*. A homologue of Sly1 was also found in H99 (CNAG_05933) which is annotated as a hypothetical protein and identical at 36% to ScSly1 from *S. cerevisiae*. Likewise, a homologue of ScSec1, the syntaxin-binding protein 1 (CNAG_07800), was found with 28% identity in H99. The roles of CNAG_05933 and CNAG_07800 have yet to be characterized.

Overall, this study provided new insights into the machinery that mediates essential nutrient uptake, trafficking of material between organelles and the relevance of endocytosis to the pathogenicity of *C. neoformans.* Many questions remain regarding the roles of additional trafficking components and the mechanisms of delivery to target organelles, particularly in the context of iron and heme delivery to mitochondria.

## MATERIAL AND METHODS

### Strains and media

The serotype A strain H99 of *Cryptococcus neoformans* var. *grubii* was used for all experiments and was maintained on YPD medium (1% yeast extract, 2% peptone, 2% dextrose, 2% agar). The Cfo1-GFP strain was constructed as described previously (9). Growth under low iron conditions was performed in yeast nitrogen base (YNB) without amino acids media plus 2% dextrose at pH 7.0 with the addition of the iron chelator bathophenanthroline disulfonate (100-150μM BPS) (YNB-BPS). Defined low-iron media (LIM) was prepared as described (52) with the addition of 20 mM HEPES and 22mN NaHCO_2_. Mammalian iron sources such as human hemoglobin (2μg mL^−1^), bovine heme (10-100μM), human holo-transferrin (50μg mL^−1^), and sheep blood (0.05%), as well as ferric chloride (FeCl_3_) (10-100μM) were added to cultures. Different compounds were added at the following concentrations: 500nM *N*-Ethylmaleimide (NEM), 20μg mL^−^ 1 brefeldin A (BFA), 500μg mL^−1^ monensin, 150ng mL^−1^ tunicamycin (TNC), 25μM aminodarone (AMD), 100μg mL^−1^ cyclosporine A (CSA), 500ng mL^−1^ tacrolimus (FK506), 5mM or 50mM ethylene glycol-bis(2-aminoethylether)-*N*,*N*,*N*′,*N*′-tetraacetic acid (EGTA), 50μg mL^−1^ rotenone, 1mM malonic acid, 10 mM salicylhydroxamic acid (SHAM). 2μg mL^−1^ antimycin A, 5mM potassium cyanide (KCN), 0.01% hydrogen peroxide (H_2_O_2_), 50μM plumbagin, 5μg mL^−1^ menadione, 1 mg mL^−1^ calcofluor white (CFW), 0.01% sodium dodecyl sulfate (SDS), 0.5mg mL^−1^ caffeine, 1.5M sodium chloride (NaCl), 10 μg mL^−1^ caspofungin, 1.6mM quinacrine, 6mM chlroquine, 10μg mL^−1^ fluconazole and 50ng mL^−1^ miconazole. All chemicals were obtained from Sigma-Aldrich.

### Construction of deletion mutants and complemented strains

All deletion mutants were constructed by homologous recombination using gene specific knockout cassettes, which were amplified by three-step overlapping PCR (53) with the primers listed in S1 Table. The resistance marker for nourseothricin (NAT) was amplified by PCR using the primers Cassette F and Cassette R and the plasmid pCH233 as a template. The gene-specific knockout primers 1 and 2, and 3 and 4 were used to amplify the flanking sequences of their respective genes; and primers 5 and 6 were used to amplify the gene-specific deletion construct containing the resistance marker. All constructs for deletions were introduced into the H99 wild-type (WT) and Cfo1-GFP strains by biolistic transformation, as described previously (54). Complementation of the *vps45*Δ mutants was performed by cloning a PCR product of *VPS45* into the integrative Safe Haven vector pSDMA58. The plasmid construct was linearized with PacI and introduced into the *vps45* and Cfo1-GFP strains by biolistic transformation. Multiplex PCR was performed on gnomic DNA of hygromycin-resistant colonies using primers UQ1768, UQ2962, UQ2963 and UQ3348, as described (55). The GFP coding sequence was added to the pSDMA58-VPS45 plasmid using fast cloning techniques (56). The GFP sequence was amplified from pHD58 plasmid whereas the GFP sequence flanked by the *GAL7* terminator was introduced into pSDMA58. The plasmid construct was linearized with PacI and introduced into the *vps45* strain by biolistic transformation. PCR for deletion constructs was performed using ExTAQ polymerase (TaKaRa Bio Inc) and the Phusion High Fidelity DNA Polymerase (New England Biolabs, USA) was used for complementation and fast cloning.

### Growth assays

To assess growth on solid media, 10-fold serial dilutions of cells were spotted on YPD agar plates containing the compounds as indicated in the text. Plates were incubated at 30°C and 37°C for 2-5 days before being photographed. For growth assays in liquid and solid YNB-BPS media, cells were pre-grown in YNB + 150 μM BPS for 48h at 30°C, washed 3 times with iron-free water (chelex-treated water) and counted. 1×10^5^ cells mL^−1^ were inoculated into YNB-BPS + iron sources for liquid assays and 10-fold serial dilutions (started at 1×10^6^ cells mL^−1^) were spotted onto plates of solid media. Flasks and plates were incubated at 30°C and 37°C and optical density at 600 nm was measured after 4h and every 24h using a spectrophotometer (BECKMAN DU^®^530) for liquid assays, whereas plates were photographed after 2-4 days.

### Microscopy

Staining of cells with the lipophilic dye FM4-64 [N-(3-triethylammoniumpropyl)-4-(6-(4-(diethylamino) phenyl) hexatrienyl) pyridinium dibromide] (T-3166; Invitrogen, Ontario, Canada) was performed to observe internalized vesicle trafficking and vacuolar membrane. The cells were stained with 5 μM for FM4-64 in PBS. The cells were incubated for an additional 90 min at 30°C and 37°C and pictures were taken every 15 min using an Olympus Fluoview FV1000 laser scanning confocal system. Colocalization analyses were performed on images of whole cells using the ImageJ coloc2 test. Positive correlations between VPS45-GFP and subcellular region were determined by Pearson’s R value (0.15-0.65) and Cortes P-value (0.95-1.00). A total of 20-35 cells were analysed per condition.

### Cell wall and mitochondria staining and flow cytometry

Cells were grown overnight at 30°C and 37°C in YPD, washed in PBS, and resuspended at 10^7^ mL^−1^ for staining as follows (all manipulations at RT). For Eosin Y (Sigma), cells were resuspended in McIlvaines buffer, pH 6.0, and stained with 250μg mL^−1^ of the dye. For Pontamine (Pontamine fast scarlet 4B, Bayer Corp., Robinson, PA), cells were stained in PBS with 100μg mL^−1^. For Concanavalin A-FITC (Sigma-Aldrich), cells were stained with 30μg mL^−1^ in Hepes-buffered saline, pH 7.0, containing 10mM each MgCl_2_ and CaCl_2_. Cells were stained for 15 min at room temperature, and then washed three times in appropriate buffers. Cells were then resuspended in PBS with 10mM NaN3 and analyzed by flow cytometry on a BD FACSCalibur™. The acidic pH sensor carboxy-DCFDA (5-(and-6-)-Carboxy-2’7’-dichlorofluorescein diacetate) (ThermoFisher Scientific) was used to assess intracellular acidification. Overnight cultures were stained with 50μM carboxy-DCFDA and incubated for 30 min at 30°C and 37°C. Cells were spun down and resuspended in 10mM NaN_3_ in PBS. 50, 000 cells were analyzed by flow cytometry on a BD LSR II-561. JC-1 (ThermoFisher Scientific was used to stain mitochondria and assess their membrane potential (MMP). Overnight cultures were stained with 5μM JC-1 for 30 min at 30°C and 37°C, washed 3 times, resuspended in media, and analyzed by flow cytometry on a BD LSR II-561. Flow analysis was performed using FlowJo software.

### Capsule formation

Capsule formation was examined by differential interference contrast microscopy on an Axio imager M.2 microscope (Zeiss) with magnification X1000 after incubation for 48 h at 30°C and 37°C in defined LIM and staining with India ink. Capsule measurements were performed using ImageJ. Capsule shedding from cells was examined with a blot assay using the anti-GXM 18B7 antibody as described (41).

### Intracellular iron and calcium concentration

Cellular iron and calcium contents were measured by inductively coupled plasma mass spectrometry (ICP-MS). Briefly, cells were pre-grown iron starved cells inoculated for 48h in YNB + 150 μM BPS °±100μM FeCl_3_ at 30°C or 37°C, washed 3 time with metal-chelated PBS, and lyophilized. The lyophilized cells were digested with 5 mL of HNO_3_ and 3 mL of H_2_O_2_ using a microwave digestion system (START D). Total iron and calcium contents were analyzed using an OPTIMA 5300 DV (PerkinElmer) system.

### Macrophage assay

Macrophage infections were performed as described previously (57). Briefly, macrophage-like J774.A1 cells were grown to 80% confluence in DMEM supplemented with 10% fetal bovine serum and 2mM L-glutamine at 37°C with 5% CO_2_. Macrophages were stimulated 2 h prior to infection with 150ng mL^−1^ phorbol myristate acetate (PMA). Fungal cells were grown in YPD overnight and PBS-washed cells were opsonized inDMEM with 0.5 μg mL^−1^ of the monoclonal antibody 18B7 for 30 min at 37°C. Stimulated macrophages were infected with 2×10^5^ opsonized fungal cells (MOI 1:1) for 2 and 24 h at 37°C with 5% CO_2_. Macrophages containing internalized cryptococci cells were washed thoroughly 3 times with PBS and then lysed with sterile water for 30 min. Lysate dilutions were plated on YPD agar and incubated at 30°C for 48 hrs, at which time the resulting CFUs were counted. Statistical significance of intracellular survival was determined by two-tailed unpaired t-tests (GraphPad Prism 7 for Windows, GraphPad Software, San Diego, CA).

### Virulence assays in mice

The virulence of the WT strain H99, the vps45Δ-34 mutant and the complemented strain *vps45*Δ::*VPS45* (52) was assessed using female BALB/c (4 to 6 weeks old) from Charles River Laboratories (Ontario, Canada). A mice survival assay with assessment of fungal burden at the humane endpoint was executed. Fungal cells were grown in 5mL of YPD at 30°C and washed twice with PBS (Invitrogen). Mice were anesthetized intraperitoneally with ketamine (80 mg kg^−1^) and xylazine (5.5 mg kg^−1^) and suspended on a silk thread by the superior incisors. A suspension of 5×10^4^ cells in 50μL was slowly inoculated into the nares of the mice. The health status of the mice was monitored daily post-inoculation and mice reaching the humane endpoint were euthanized by CO_2_ anoxia. Fungal burden of organs (lungs, brain, liver, spleen, and kidney) and cardiac blood was assessed. The organs and blood were aseptically removed. Blood was retrieved from the heart using sterile syringes pre-rinsed with 500 units of heparin. Organs were weighed, and homogenized using a Retsch MM301 mixer mill. The samples were serial diluted, plated on YPD containing chloramphenicol (30μg mL^−1^) and incubated at 30°C for 2 days; CFUs were then counted. Statistical analyses of survival differences in mice were performed with the log rank test and a two-tailed unpaired Mann-Whitney test was used to assess the fungal load (GraphPad Prism 7 for Windows, GraphPad Software, San Diego, CA).

### Ethics Statement

This study was carried out in strict accordance with the guidelines of the Canadian Council on Animal Care. The protocol for the virulence assays employing mice (protocol A17-0117) was approved by the University of British Columbia Committee on Animal Care.

## ACKNOWLEDGEMENTS

The authors would like to thank Dr. Arturo Casadevall for the 18b7 antibody, Dr.James A. Fraser for the safe haven plasmid pSDMA58, Dr. Eddy Sánchez-León for assistance with confocal microscopy, Dr. Hao Ding for the generation of plasmid pHD58, and Arif A. Arif for his help with flow cytometry analysis.

## SUPPORTING INFORMATION

**S1 Table: Primers used in this study**

**S1 Fig.:** Southern blot of genomic DNA for the WT strain H99, *vps45* mutants and complemented strains. Extracted DNA was digested with BglI and genomic hybridization to detect the *VPS45* locus yielded fragments of 4329 bp in the WT strain, 1582 bp in the deletion mutants, and 4478 bp and 1582 bp in complemented strains.

**S2 Fig.: Intracellular membrane staining with the lipophilic dye FM4-64.** WT, *vps45* mutant, complemented (A), and GFP-tagged (B) strains were grown for 24h in YNB-BPS at 37°.. 1×10^6^ cells/mL were inoculated in YNB-BPS + 100μM FeCl_3_, stained with 5 μM FM4-64 and transferred in a chamber slide where the cells were maintained at 37°C Confocal microscopy images were taken every 15 min for 90 minutes. Colocalization analyses using ImageJ coloc2 test revealed a positive correlation between VPS45-GFP and endocytic membranes as determined by Pearson’s R value (0.15-0.45) and Cortes P-value (0.99-1.00).

**S3 Fig.: Restoration of *vps45* mutant phenotypes by the fusion protein Vps45-GFP expressed from the safe haven location.** The WT, mutants and complemented strains (*VPS45* or *VPS45-GFP*) were grown on YPD in the presence of 10μg mL^−1^ fluconazole, 25 μM aminodarone and 6mM chloroquine. Cells were pre-cultured in YPD overnight at 30°C, serial diluted, and 5μL were spotted onto YPD plates containing the indicated drugs. Plates were incubated at 30°C or 37°C for 2 days.

**S4 Fig.: A) Localization of Cfo1-GFP in the absence of *VPS45* at 37°C.** Strains (WT, *vps45Δ* and complement) containing a *CFO1-GFP* construct were cultured overnight in YPD, washed 3 times and counted. 1×10^6^ cells/mL were inoculated in YNB-BPS and incubated at 37°C for 1h. Then cells were stained with 5 μM FM4-64, transferred in a chamber slide and maintained at 37°C. Confocal images were taken every 15 minutes for 2h. B) Localization of Cfo1-GFP after 24h incubation in YNB, YNB + 150μM BPS, and YNB-BPS + 100μM FeCl_3_ at 37°C. The GFP label indicates evaluation of the green fluorescent protein and DIC indicates differential interference contrast microscopy. C) Visualization of full-length Cfo1-GFP by Western Blot in the WT strain (1), the *vps45* mutant (2) and the complemented strain (3) cells incubated for 3h in YNB+150μM BPS at 30°C or 37°C. Western Blot on Cfo1-GFP using the mouse monoclonal GFP antibody (B-2) (Santa Cruz Biotechnology). Cfo1 is expected to be glycosylated like its homologue Fet3p in *S. cerevisiae* and this would explain the observed slower migration of the band (~140 kDa) versus the predicted size of the fusion (~100 kDa; Cfo1: 71.1 kDa and GFP: 27 kDa) (59). D) Blue silver staining of the SDS-PAGE gel with ~10 ug of total cell protein extracted by bead beating and sonication from cells incubated for 3h in YNB + 150μM BPS at 30°C or 37°C.

**S5 Fig.: Influence of Vps45 on mitochondrial function, ROS sensitivity and mitochondrial membrane potential.** Disruption of *VPS45* did not influence mitochondria morphology in iron-chelated conditions. Confocal images of mitochondria stained with 500nM mitotracker were taken of the WT, mutant and complemented strains inoculated for 24h in YNB ° 150μM BPS at 30°C (A) and 37°C (B). C). The WT, mutants and complemented strains were grown in the presence of inhibitors of the mitochondrial electron transport chain and ROS. Cells were pre-cultured in YPD overnight at 30°C, serial diluted, and 5μL were spotted onto YPD plates containing A) 75μg mL^−1^ rotenone, 1mM malonic acid, 2μg mL^−1^ oligomycin A, B) 500μM paraquat, 50μM plumbagin, 50μM diphenyleneiodonium chloride (dpi) or 5μg mL^−1^ menadione. Plates were incubated at 30°C and 37°C for 2 days. D) Example of flow cytometry assessment of mitochondrial membrane potential with the JC-1 dye. The percentages of cells with polarized, mixed and depolarized mitochondria membrane potential were determined by gating the fluorescence of unstained single cells population to stained single cells population.

**S6 Fig.: Growth and survival of WT, *vps45* mutants and complemented strains in defined low iron media** (33). Cells were pre-grown in defined LIM for 48h at 30°C, washed and counted. 5×10^5^ cells/mL were inoculated into LIM (A and C), LIM + 100 μM FeCl_3_ (B and D) and incubated at 30°C (A and B) or 37°C (C and D) for 48h.

